# Identification of a CCG-enriched expanded allele in DM1 patients using Amplification-free long-read sequencing

**DOI:** 10.1101/2022.02.22.481438

**Authors:** Yu-Chi Tsai, Laure de Pontual, Cheryl Heiner, Tanya Stojkovic, Denis Furling, Guillaume Bassez, Geneviève Gourdon, Stéphanie Tomé

## Abstract

Myotonic dystrophy type 1 (DM1) exhibits highly heterogeneous clinical manifestations caused by an unstable CTG repeat expansion reaching up to 4,000 CTG. The clinical variability depends on CTG repeat number, CNG repeat interruptions and somatic mosaicism. Currently, none of these factors are simultaneously and accurately determined due to the limitations of gold standard methods used in clinical and research laboratories. An amplicon method for targeting DM1 locus using Single-Molecule Real-Time sequencing was recently developed to accurately analyze expanded alleles. However, amplicon-based sequencing still depends on PCR and the inherent bias towards preferential amplification of smaller repeats can be problematic in DM1. Thus, an amplification-free long-read sequencing method was developed using the CRISPR/Cas9 technology in DM1. This method was used to sequence the DM1 locus in patients with CTG repeat expansion ranging from 130 to > 1000 CTG. We showed that elimination of PCR amplification improves the accuracy of measurement of inherited repeat number and somatic repeat variations, two important key factors in the DM1 severity and age at onset. For the first time, an expansion composed of over 85% CCG repeats was identified using this innovative method in a DM1 family with an atypical clinical profile. No-Amplification targeted sequencing represents a promising method that can overcome research and diagnosis shortcomings, with translational implications for clinical and genetic counseling in DM1.

## Introduction

Short tandem repeats are an important source of genetic variation and phenotypic variability in disease and health, that are not always well characterized and understood due to their complexities. Among these repeated elements, unstable repeat expansions are associated with more than twenty diseases including the complex and variable myotonic dystrophy type 1 (DM1) disorder^1^. DM1 is caused by an unstable CTG repeat expansion in the 3’UTR of the *dystrophia myotonica protein kinase* (*DMPK*) gene, which usually increases across generations and over time in somatic tissues^2,3^. In DM1, longer expanded alleles are usually associated with a worsening of clinical severity and an earlier age of onset^4^. This anticipation phenomenon is particularly obvious in DM1^5^. DM1 is mainly characterized by an unusually broad clinical spectrum of symptoms divided into five distinct clinical forms ranging from late onset to the congenital forms which are often associated with the largest size of inherited disease-associated allele^4^. Facial dysmorphisms, muscle weakness and cognitive impairment are more frequent symptoms at an earlier onset, while cardiac defects and cataracts are more common in DM1 patients with later forms of the disease^4^.

In DM1, it is laborious to diagnose patients and classify them into distinct clinical categories based exclusively on mutation status and size of CTG repeats when known, for several reasons. First, the long-expanded allele and the precise number of CTG repeats are complicated to identify and measure by conventional methods, particularly for the larger repeat expansions^6,7^. Second, DM1 patients exhibit high clinical and genetic variability which cannot be exclusively explained by the size of the CTG repeats^4^. Third, additional factors such as somatic mosaicism are important disease modifiers contributing to the high genotype and phenotype variability observed in DM1. Somatic mosaicism biased towards expansions contributes to the progressive nature of the various DM1 symptoms and also to the variation in the age of onset^8–11^. Forth, the majority of DM1 patients inherit pure CTG repeat expansion. However, more than 8% of known DM1 patients carry interruptions that vary in type (CCG, CAG, CTC and CGG) and number between families and also among individuals of the same family^12^. Interruptions are frequently associated with intergenerational contractions and stabilization of the CTG repeat as well as milder DM1 symptoms and / or additional symptoms^11,13–22^. Fifth, single-nucleotide polymorphisms in the DNA mismatch repair gene *MSH3*, required for maintenance of genomic integrity, have been shown to reduce levels of somatic mosaicism and are associated with delayed onset in DM1 patients^23,24^. The contribution of CTG repeat length, somatic mosaicism, structural variants and/or modifier genes on the DM1 genotype and phenotype remains poorly understood due to technical difficulties in analyzing them.

Currently, the molecular diagnostic procedure used in DM1 genetic testing is divided into two steps^6,7^. The first is to detect or rule out possible DM1 expansions using PCR and fragment-length analysis (Table 1). This method cannot differentiate between individuals homozygous for a normal allele and individuals whose CTG repeat expansion could not be amplified by PCR. A second step is therefore necessary to identify large expanded alleles and estimate the approximate size using Southern blotting. All of these methods are time-consuming procedures that yield no information on somatic mosaicism and structural variants of expansion, two important prognostic parameters. Furthermore, triplet-repeat primed PCR (TP-PCR) is also used to directly detect the presence of an expanded allele^25^. However, no size repeat can be estimated by this method, which leads to a loss of information necessary for carrying out genetic counseling. To date, TP-PCR is the only diagnostic method which makes it possible to reveal variant repeats in the first 100bp within the 5’ and 3’ end of CTG array. However, this method is unable to identify interruptions at the middle of large repeat expansions of more than 200-300 CTG repeats and to give the exact number and type of interruptions, regardless of the size of the repeats. Diagnostic laboratories generally use several methodological approaches to detect CTG repeat expansion and roughly estimate the size of the repeat which leads to long diagnostic delays that can exceed 3 months. Unfortunately, no diagnostic method can estimate somatic mosaicism which is an important parameter to predict disease progression^8–11^. Today, the most accurate method to assess the inherited CTG repeat expansion size and the level of somatic mosaicism is the Small-Pool PCR (SP-PCR)^9,26^. SP-PCR, PCR on successive DNA dilutions, is a time-consuming and fastidious method, exclusively used in the research laboratories. The PCR amplification bias towards shorter repeats and the difficulties inherent to the amplification of the GC rich DM1 locus by this method remain a concern, which needs to be addressed by a new widely available method. In addition, neither the SP-PCR nor Southern blotting can detect CNG interruptions within the repeats, which can lead to inaccurate genetic analysis and genotype-phenotype associations. Currently, molecular diagnostic tests used in diagnostic laboratories to analyze the DM1 locus may induce loss of information and bias in the interpretation of genetic data, especially when patients carry large CTG repeats. It is therefore essential to accurately measure the number of CTG repeats, structural variants and somatic mosaicism in DM1 patients using a fast and accurate method to provide better diagnosis and prognosis with minimal delay. Pacific Biosciences’ (PacBio) targeted single-molecule long-read real-time sequencing from PCR products was recently described to analyze the DM1 locus in patients^16,27^. This method allows accurate counting of repeat numbers in large expanded alleles, and can simultaneously evaluate the degree of somatic mosaicism and identify interruptions^27^. However, amplicon-based sequencing relies on PCR and is still subjected to the inherent preferential amplification of smaller repeats, which introduces a confounding and undesirable bias in repeat sizing. To overcome this limitation, we developed a robust amplification free-targeted long-read sequencing for DM1. Using this method, measurement of CTG repeat length and somatic mosaicism is improved in DM1 patient samples. The second advantage of this method is that CNG interruptions are easily detected regardless of the size of the CTG repeat expansions. Strikingly and for the first time, we identified triplet repeat expanded alleles composed of more than 85% of CCG in DM1 patients for whom amplification of the triplet repeat expansion failed by PCR and TP-PCR.

**Table 1:** Genetic testing strategy for DM1.

| Information | PCR | Southern blot | TP-PCR |
| --- | --- | --- | --- |
| Mutation Status | √ | √ | √ |
| CTG repeat size<br><200-300 | √ | √ | √ |
| Large CTG repeat<br>size | x | √ | x |
| Structural variants | x | x | x |
| Somatic mosaicism | x | x | x |

## Materials and Methods

### DM1 patient genomic DNA samples

DM1 participants were recruited by the Genetics Department of Nantes Hospital, the Genetics Department of the Necker-Enfants Malades Hospital, the DM-Scope registry and the Neuromuscular Disease Reference Center of Pitié-Salpêtrière Hospital in France. Written informed consent was obtained from all participants. Genomic DNA samples were initially genotyped for DM1 using conventional PCR, TP-PCR, or Southern Blot^6,7^. DNA was extracted in diagnostic laboratories. High molecular weight (HMW) DNA was extracted from immortalized lymphoblastoid cells by the Monarch® Genomic DNA Purification Kit from New England Biolabs.

### Design of Guide RNAs for cas9 digestion

Human Hg19 reference sequences surrounding the DM1 repeat region were used to design Cas9 CRISPR RNA (crRNA) sequences. Candidate target sequences were generated using the GPP sgRNA Designer webtool on the Broad Institute website (https://portals.broadinstitute.org/gpp/public/analysis-tools/sgrna-design). Two upstream and three downstream top ranking candidate sequences which could generate approximately 2kb target fragments were custom synthesized and tested in targeted sequencing experiments with HEK293 genomic DNA. The combination of upstream and downstream crRNAs that produced the highest sequencing yield was selected for this study. The final crRNA sequences are the DM1-L1-crRNA sequence: 5’-CCCCATCGGGACAACGCAGA-3’ and the DM1-R1-crRNA sequence: 5’-GGGCGTGTATAGACACCTGG-3’ which generates a target capture region of 2361 bp.

### SMRT bell library preparation

Preparation of the SMRTbell® library was performed according to the *Procedure and Checklist-No-Amp Targeted Sequencing utilizing the CRISPR-Cas9 system*, available on the PacBio website (https://www.pacb.com/wp-content/uploads/Procedure-Checklist-No-Amp-Targeted-Sequencing-Utilizing-the-CRISPR-Cas9-System.pdf). Approximately 3-5μg of native genomic DNA were dephosphorylated with 0.05U/μl of shrimp alkaline phosphatase (rSAP) (New England BioLabs) to reduce the amount of off-target molecules in the final SMRTbell® library. Guide RNAs (gRNAs) were formed by annealing crRNAs (Integrated DNA Technologies) to tracrRNAs (Integrated DNA Technologies) at a 1:1 ratio and 5μM concentration. The sequences of each crRNA targeting DM1 and various control loci used in the experiments are shown in Table 2. The gRNA-Cas9 complex was prepared by incubating 400nM gRNA with 400nM Cas9 nuclease (New England Biolabs) for 10 min at 37°C. Dephosphorylated genomic DNAs were digested with the gRNA-Cas9 complex for 1 hour at 37°C. DM1 and control gRNA were multiplexed in the same digestion reaction. Cas9 digestion products were purified using a 0.45X volume of AMPure PB beads (Pacific Biosciences). Barcoded adapters recommended by PacBio were ligated to purified Cas9 digestion products using 0.4μM barcode adapters and 0.9U/μl T4 DNA ligase (Thermo Fisher Scientific) to form a library of symmetric SMRTbell template molecules. The SMRTbell® library was purified using a 0.45X volume of AMPure® PB beads (Pacific Biosciences, Menlo Park, CA, USA). Failed ligation products and gDNA fragments were removed by nuclease treatment using 1.2U/ul of Exonuclease III (New England BioLabs) and an Enzyme Clean Up Kit (Pacific Biosciences, Menlo Park, CA, USA) for 2 hours at 37°C. After nuclease treatment, the SMRTbell® library was incubated with 41μg/mL SOLu-Trypsin (Sigma-Aldrich) for 20 minutes at 37°C. The nuclease-treated SMRTbell® library was purified using a 0.45X volume of AMPure® PB beads for the first purification and a 0.42X volume of AMPure® PB beads for the second purification. Finally, the purified SMRTbell library was resuspended in 6.3μl of elution buffer for targeted SMRT® sequencing.

**Table 2:**
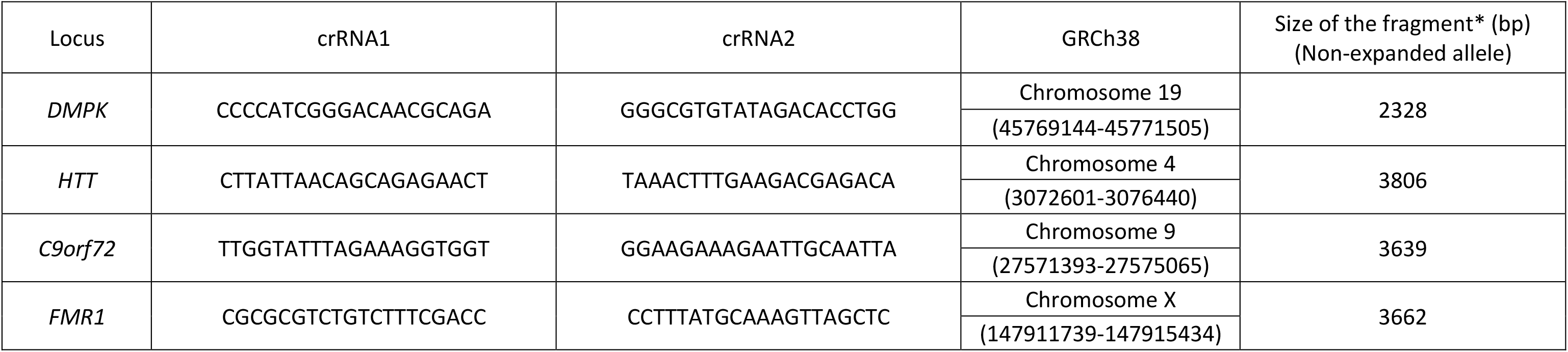
crRNA and genomic coordinate for *DMPK* and control loci (* Fragment size after CRISPR/Cas9 digestion)

### Targeted SMRT sequencing

Following *the Procedure and Checklist-No-Amp Targeted Sequencing Utilizing the CRISPR-Cas9 System* (https://www.pacb.com/wp-content/uploads/Procedure-Checklist-No-Amp-Targeted-Sequencing-Utilizing-the-CRISPR-Cas9-System.pdf), final SMRTbell® libraries were annealed with Sequencing Primer v4 from the No-Amp Accessory Kit (Pacific Biosciences, Menlo Park, CA, USA) and bound with Sequel® DNASequencing Polymerase 3.0 in the Sequel Binding Kit 3.0 (Pacific Biosciences, Menlo Park, CA, USA). Polymerase-bound SMRTbell complexes were purified with 0.6X AMPure® PB beads and sequenced on a PacBio Sequel System using Sequel Sequencing Kit 3.0 and a customized setting with 4 hours of complex immobilization and 20-hour movie collection.

### Sequencing data and repeat sequence analysis

Sequencing data repeat sequence analysis were performed following the Analysis Procedure – No-Amp Data Preparation and Repeat Analysis available on the PacBio website (https://www.pacb.com/wp-content/uploads/Analysis-Procedure-No-Amp-Data-Preparation-and-Repeat-Analysis.pdf). Barcode demultiplexed high accuracy long reads (HiFi reads) were first generated using the PacBio SMRT Link software v8.0 or later version. HiFi reads provide accuracy of >99.9%. HiFi reads corresponding to each sample were mapped to GRCh38 human reference sequence, allowing on-target read identification and visual inspection of sequencing reads using Integrative Genomics Viewer^28^. Repeat sequence analysis was then performed using the RepeatAnalysisTools and instruction available through GitHub (https://github.com/PacificBiosciences/apps-scripts/tree/master/RepeatAnalysisTools) to generate various repeat sequence statistics,

### Amplicon long-read targeted sequencing

DM1 locus was characterized in DM1 individuals 11 and 18 according to the method described in Mangin et *al*.^27^ Briefly, normal and expanded CTG repeat alleles were amplified by PCR using barcoded ST300-F (5’-GAACTGTCTTCGACTCCGGG-3’) and ST300-R (5’-GCACTTTGCGAACCAACGAT-3’) primers. In order to enrich the SMRTbell® library with the expanded allele, the PCR products were purified using a 0.45-0.5X volume of AMPure® PB beads (Pacific Biosciences, Menlo Park, CA, USA) prior to SMRT sequencing. SMRTbell® libraries were prepared using the SMRTbell Express Template Prep Kit 2.0 and sequenced using a sequel system.

### Small Pool PCR

Small pool PCR (SP-PCR) amplifications and PCR product electrophoretic analyses were performed according to Gomes-Pereira et *al*. ^29^. Briefly, blood DNA samples were digested with *HindIII* enzyme and SP-PCR was performed using DM-C (5’-AACGGGGCTCGAAGGGTCCT-3’) and DM-BR (5’-CGTGGAGGATGGAACACGGAC-3’) primers^26^. The conditions of PCR were as follows: denaturation at 96°C for 5 min followed by 30 cycles at 96°C for 45s, 68°C for 30s and at 70°C for 3 min with a chase of 1 min at 68°C and 10 min at 72°C. SP-PCR products were loaded on a 1% agarose gel to compare instability between DM1 patients 11 and 18 and ran at 300V for 30 min followed by 16 hours at 160V. The PCR products were then transferred to GeneScreen Plus Hybridization Transfer Membrane (Perkin Elmer) and detected by hybridization using a non-radioactive method as described in Tomé et *al*.^30^.

## Results

### Amplification-free DM1 locus enrichment in DM1 patients

Amplicon-based long-read sequencing depends on PCR, which has an inherent bias towards smaller repeats. To overcome this limitation, an amplification-free long read sequencing method (No-Amp) was used to specifically characterize the DM1 locus. This method utilizes the CRISPR/Cas9 system to target and isolate the DNA fragment of interest from genomic DNA, in combination with long-read single-molecule real-time (SMRT) sequencing. This method was performed in seven DM1 patients using DNA isolated either from blood or lymphoblastoid cell lines. These patients carried different CTG repeat expansion lengths that were previously defined by Southern Blot and/or PCR. The number of generated on-target HiFi reads was sample-dependent and ranged from 114 to 733 *DMPK* reads (Table 3). The CTG repeat region was successfully captured in all samples, with a better result for samples #A3, #15969 and #15841. For those samples, DNA was extracted with a high molecular weight (HMW) DNA extraction kit (New England Biolabs). Off-target reads were observed for all samples and represent an average number of approximately 54%. No significant accumulation of off-target sequencing reads mapping to a specific genome location was observed in the samples digested exclusively with DM1 locus guide RNAs; all the off-target sequencing reads appeared to be randomly distributed across the GRCh38 reference (Figure 1). However, some specific off-target reads on chromosome 2 (chr2:116779778-116785717) and chromosome 13 (chr13:89237016-89240938) were captured in two of the three samples digested by DM1 and control locus guide RNAs (Table 2 and Figure 1). These specific off-target reads are potentially produced by off-target guide RNA sequence (DMPK-crRNA2, C9orf72-crRNA1 and C9orf72-crRNA2) matches with 2-4 mismatched bases (data not shown). Off-target as well as control reads were filtered out during the bioinformatics analysis and did not affect analysis of DM1-specific reads. For samples #L1 and #L3 carrying the largest CTG repeat sizes, number of highly accurate long reads from the expanded allele was lower than for the other samples in this study. The lower sequencing yields for these large alleles are likely the results of reduced efficiency of immobilization of DNA-polymerase complex into the SMRT cell via diffusion and lower raw reads to HiFi reads conversion efficiency due to significantly longer template lengths.

**Table 3:** HiFi reads in DM1 patients (**Includes *DMPK* and control loci reads)

| Individuals | #11 | #18 | #L1 | #L3 | #A3 | #15969 | #15841 |
| --- | --- | --- | --- | --- | --- | --- | --- |
| Tissue | Blood | Blood | Blood | Blood | Lymphoblastoid cells | Lymphoblastoid cells | Lymphoblastoid cells |
| DNA extraction | N/D | N/D | N/D | N/D | Monarch Genomic DNA Purification Kit (NEB) | Monarch Genomic DNA Purification Kit (NEB) | Monarch Genomic DNA Purification Kit (NEB) |
| HiFi reads | 841 | 1041 | 836 | 3731 | 7405 | 11201 | 4429 |
| *DMPK* locus | 452 | 352 | 332 | 114 | 569 | 733 | 597 |
| Control loci | none | none | none | none | 4522 | 3073 | 3218 |
| Expanded allele | 206 | 205 | 65 | 21 | 303 | 398 | 361 |
| Non-Expanded allele | 246 | 147 | 262 | 93 | 266 | 335 | 236 |
| % reads on-target | 53,7 | 33,8 | 39,71 | 3,06 | 68,75** | 33,98** | 86,14** |
| % reads off-target | 46,3 | 66,2 | 60,29 | 96,94 | 31,25 | 66,02 | 13,86 |

**Figure 1:**
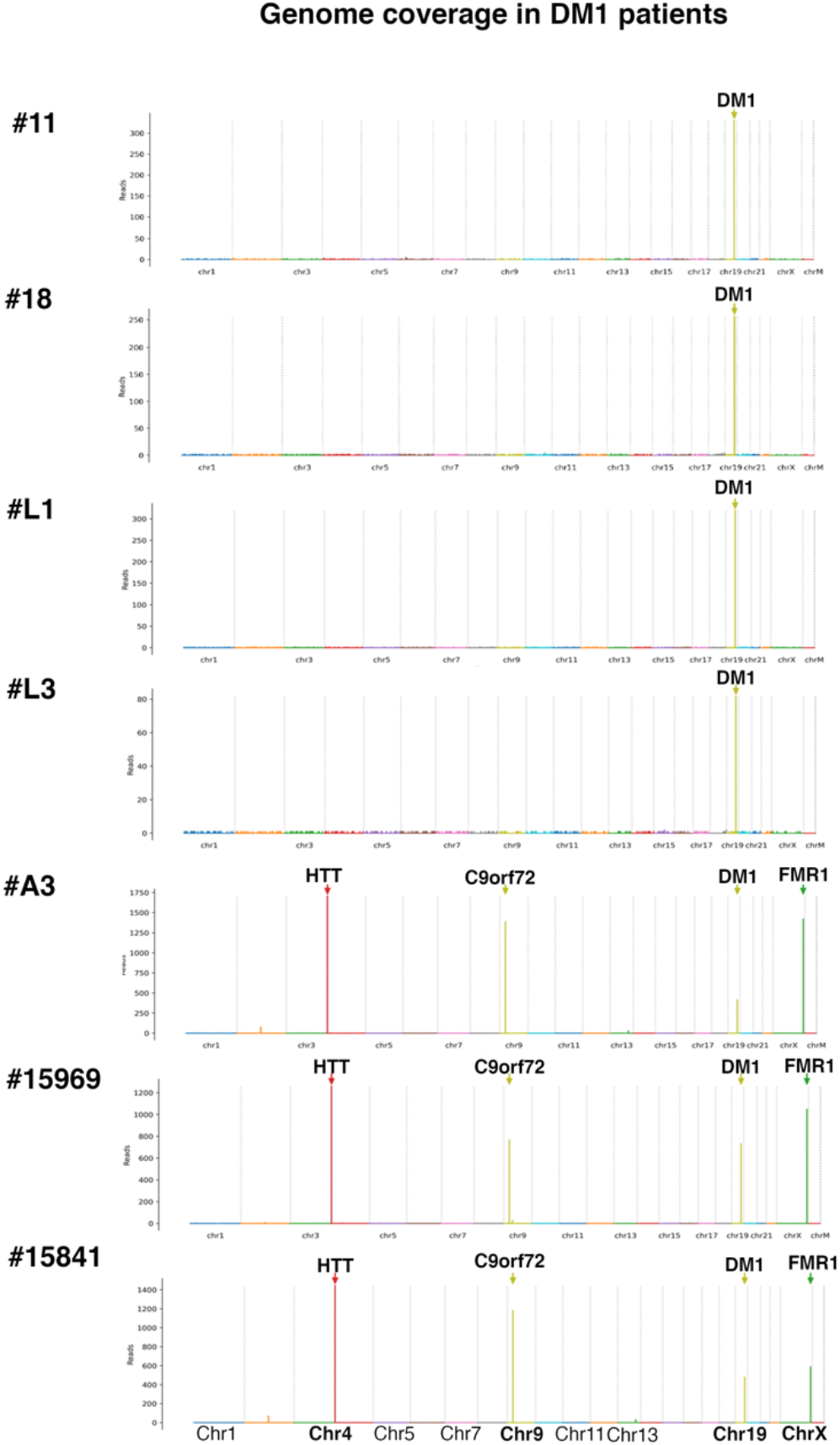
Genome wide coverage plots. Genome wide coverage plots for DM1 patients are shown in the figure. The y-axis shows the number of reads and the x-axis spans over all the chromosomes in the Hg38 genome.

### Targeted amplification-free long-read sequencing improves the accuracy of CTG repeat size measurement

This is the first time that amplification-free targeted sequencing has been described to generate long-read sequencing data analysis for the DM1 locus using the RepeatAnalysisTools developed by PacBio (see Materials and Methods for detail). By amplification-free sequencing, the number of CTG repeats was quantified in seven DM1 patients and ranged from 141 to 1378 repeats (median) (Table 4). The expanded alleles showed a median CTG repeat of 516 and 495 in patients #11 and #18, respectively, which was a similar range to that observed for amplicon target sequencing (Table 4). On the contrary, individuals #L1 and #L3 whose size was estimated to be more than 1000 CTGs in diagnosis, showed different CTG repeat medians between amplicon sequencing and no-amplification (No-Amp) targeted sequencing. The median CTG repeat size was estimated to be 563 and 927 CTG in individuals #L1 and #L3, respectively, by the amplicon sequencing experiment, and 1378 CTG and 1287 CTG by the No-Amp experiment (Table 4). These results strongly suggest that amplicon sequencing is subjected to the inherent preferential amplification of the smallest repeats, particularly in individuals #L1 and #L3 carrying the largest inherited expanded alleles. Thus, PCR may introduce a confounding and undesirable bias in repeat sizing in DM1 patients. The median CTG repeat size for the expanded allele in samples #A3 and #15969, based on No-Amp targeted sequencing, is consistent with previous Amplicon targeted sequencing data from Mangin et *al*.^27^. Interestingly, the number of repeats in patient #15841 was estimated to be 467 CNG repeats by the No-Amp method. This size estimate could not be done before by PCR because it failed several times.

**Table 4:** CTG repeat size and interruptions in DM1 patients.

| **Individuals** | **#11** | **#18** | **#L1** | **#L3** | **#A3** | **#15969** | **#15841** |
| --- | --- | --- | --- | --- | --- | --- | --- |
| **Tissue** | **Blood** | **Blood** | **Blood** | **Blood** | **Lymphoblastoid cells** | **Lymphoblastoid cells** | **Lymphoblastoid cells** |
| **Age at sample**<br>**(Years)** | **20** | **42** | **29** | **26** | **40** | **36** | **58** |
| **Median Expanded**<br>**allele**<br>**(No-Amp)** | **516** | **495** | **1378** | **1287** | **141** | **158** | **467** |
| **Median Expanded**<br>**allele**<br>**(Amplicons)** | **468** | **460** | **563** | **927** | **#N/A** | **#N/A** | **Failed** |
| **Interruptions**<br>**(No-Amp)** | **2CCG** | **4CCG** | **Pure** | **Pure** | **1CAG** | **Pure** | **78CCG** |
| **Interruptions**<br>**(Amplicons)** | **2CCG** | **4CCG** | **Pure** | **Pure** | **#N/A** | **#N/A** | **Failed** |
| **CNG repeat size**<br>**(Diag)** | **≅700** | **≅400** | **>1000** | **>1000** | **130** | **150** | **370** |

### Targeted amplification-free long-read sequencing exhibits somatic mosaicism in DM1 patients

The distribution of CNG repeats was analyzed in individuals #11 and #18 carrying less than 500 interrupted CNG repeats by targeted long-read sequencing as well as by Small-pool PCR which is usually used to estimate somatic mosaicism in DM1 patients. All these methods, including SP-PCR, showed that somatic mosaicism was lower in individual #18 than in individual #11 (Figure 2). Despite similar somatic mosaicism observed with the different methods, a PCR bias where the shortest alleles were overrepresented using long read sequencing of PCR products (Amp) is clearly observed. Interestingly, approximately 200 single molecule HiFi reads of the expanded allele appear sufficient to accurately estimate somatic mosaicism in patients by no-Amplification targeted sequencing (Figure 2). Somatic mosaicism was also observed in blood from patients #L1 and #L3 carrying more than 1000 pure CTG repeats at the DM1 locus (Figure 3A). Despite a limited number of HiFi reads obtained, no-Amplification targeted sequencing provides a better estimate of CTG repeat number distribution in patients with large expansions where PCR bias is extremely high using amplification targeted sequencing. To complement our data, we have also shown that lymphoblastoid cell lines from individuals #A3 and #15969 exhibit somatic mosaicism to varying degrees, with the highest being observed in patient #A3 (Figure 3B).

**Figure 2:**
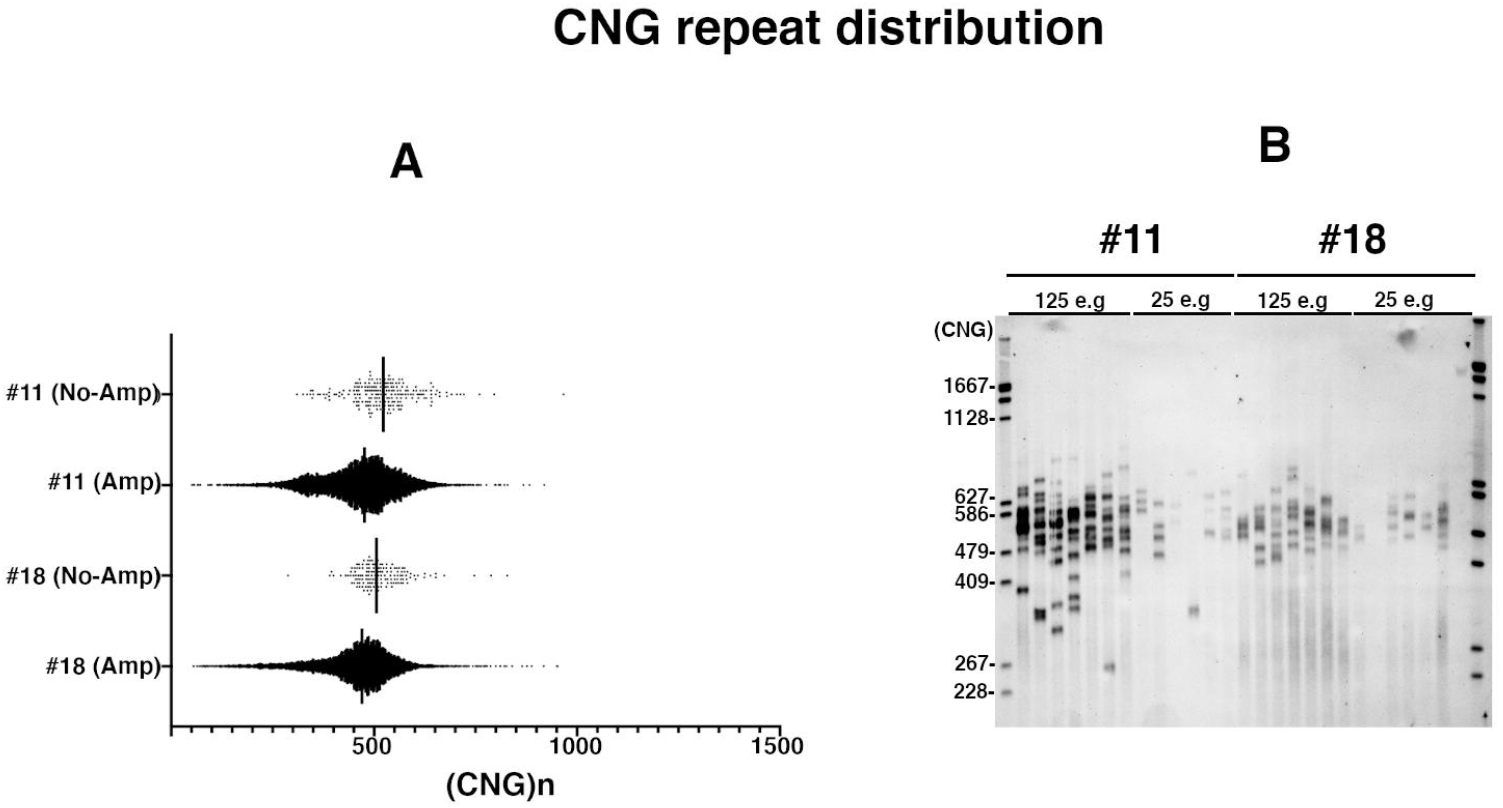
Somatic mosaicism in #11 and #18 individuals. **A-** The distribution of the size of the repeat of each HiFi read generated by PacBio algorithm. The x-axis shows the CTG repeat number. Each dot represents the CTG repeat size of one HiFi read. The black bar represents the median. 10000-30000 single molecule HiFi reads of expanded allele are analyzed in the Amplicon experiment whereas more than 200 single molecule HiFi reads of expanded allele are analyzed in the No-Amp experiment. B-Trinucleotide repeat instability estimation by Small-Pool PCR in blood samples of DM1 patients. The sizes (converted in CTG repeats) of DNA molecular weight marker III, DIG-labeled are indicated on the left of the figure. About 25-125 DNA molecules are amplified for each individual. e.g means equivalent genome.

**Figure 3:**
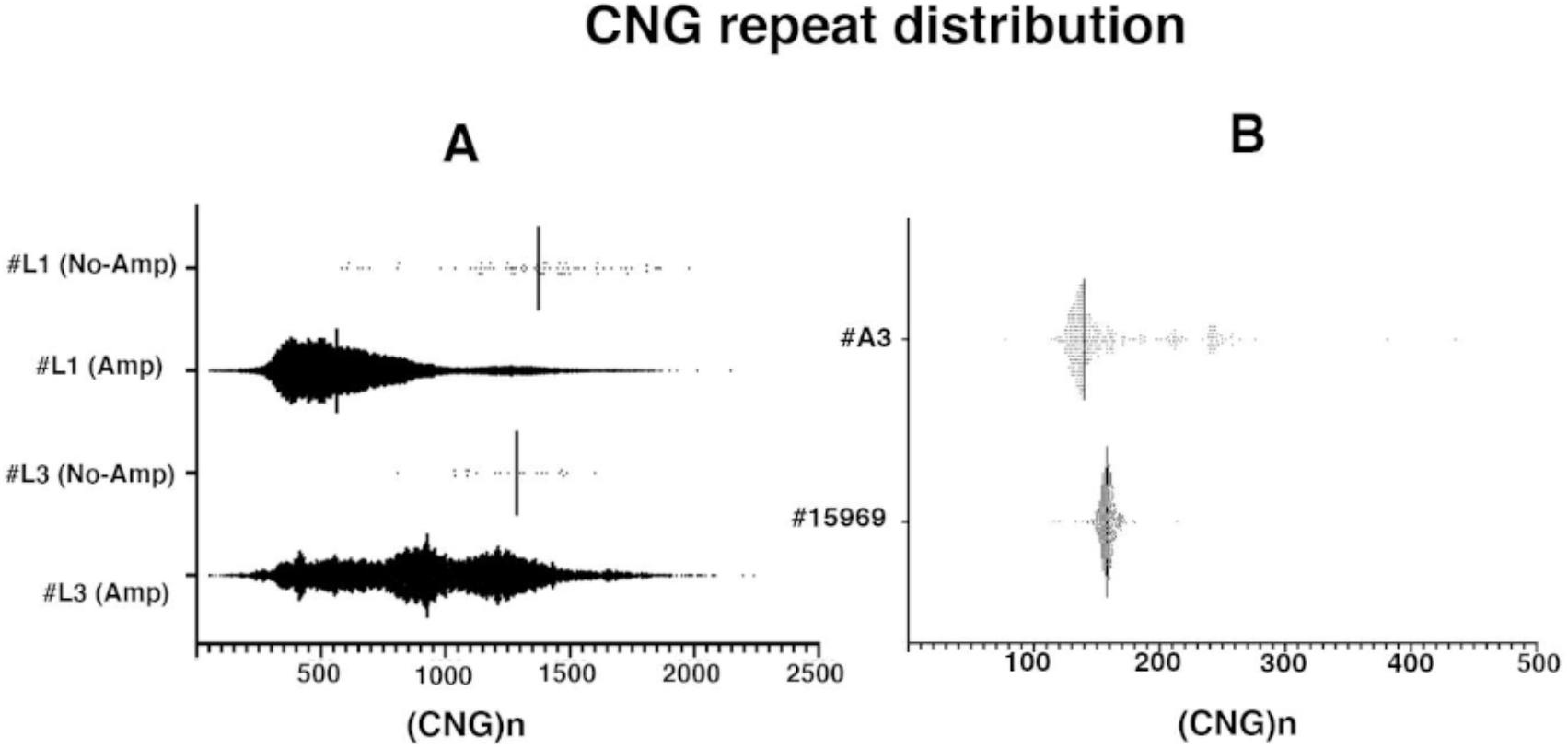
Distribution of the size of the repeat of each HiFi read generated by PacBio algorithm. A-#L1 and #L3 individuals. B-#A3 and #15969 individuals. The x-axis shows the CTG repeat number. Each dot represents the CTG repeat size of one HiFi read. The black line represents the median. 10000-30000 single molecule HiFi reads of expanded allele are analyzed in the Amplicon experiment whereas 21-398 single molecule HiFi reads of expanded allele are analyzed in the No-Amp experiment.

### Targeted amplification-free long-read sequencing detects CNG interruptions

In the literature, the interrupted expanded allele has recently been shown to be a key factor in the severity of DM1 symptoms and the age of onset of these symptoms^1,13–22^. The identification of interruptions in DM1 patients at diagnosis is therefore important to improve genetic counseling in DM1. Using the no-amplification method, we analyzed repeat expansion sequences to identify interrupted expanded alleles in DM1 patients. Samples #11 and #18 showed interruptions at the 3’ end of the CTG repeat expansion, which were also detected by TP-PCR (Figure 4 and data not shown). Two CCG interruptions and four CCG interruptions were identified in samples #11 and #18, respectively, by targeted sequencing of the DM1 locus, identification which was impossible with other gold standard methods (Figure 4). Interestingly, a single CAG repeat interruption at the 5’ end of the CTG repeat expansion was also identify in the blood-derived lymphoblastoid cell line of sample #A3 (Figure 4). In sample #15841, amplification of the expanded allele failed by PCR, making it impossible to analyze the CNG repeat expansion sequence. By targeted no-amplification sequencing, we overcame this limitation and identified approximately 78 CCG interruptions in this patient’s mutated allele located both at the 3’ end of the CTG repeat expansion and in the middle of the sequence (Figure 4).

**Figure 4:**
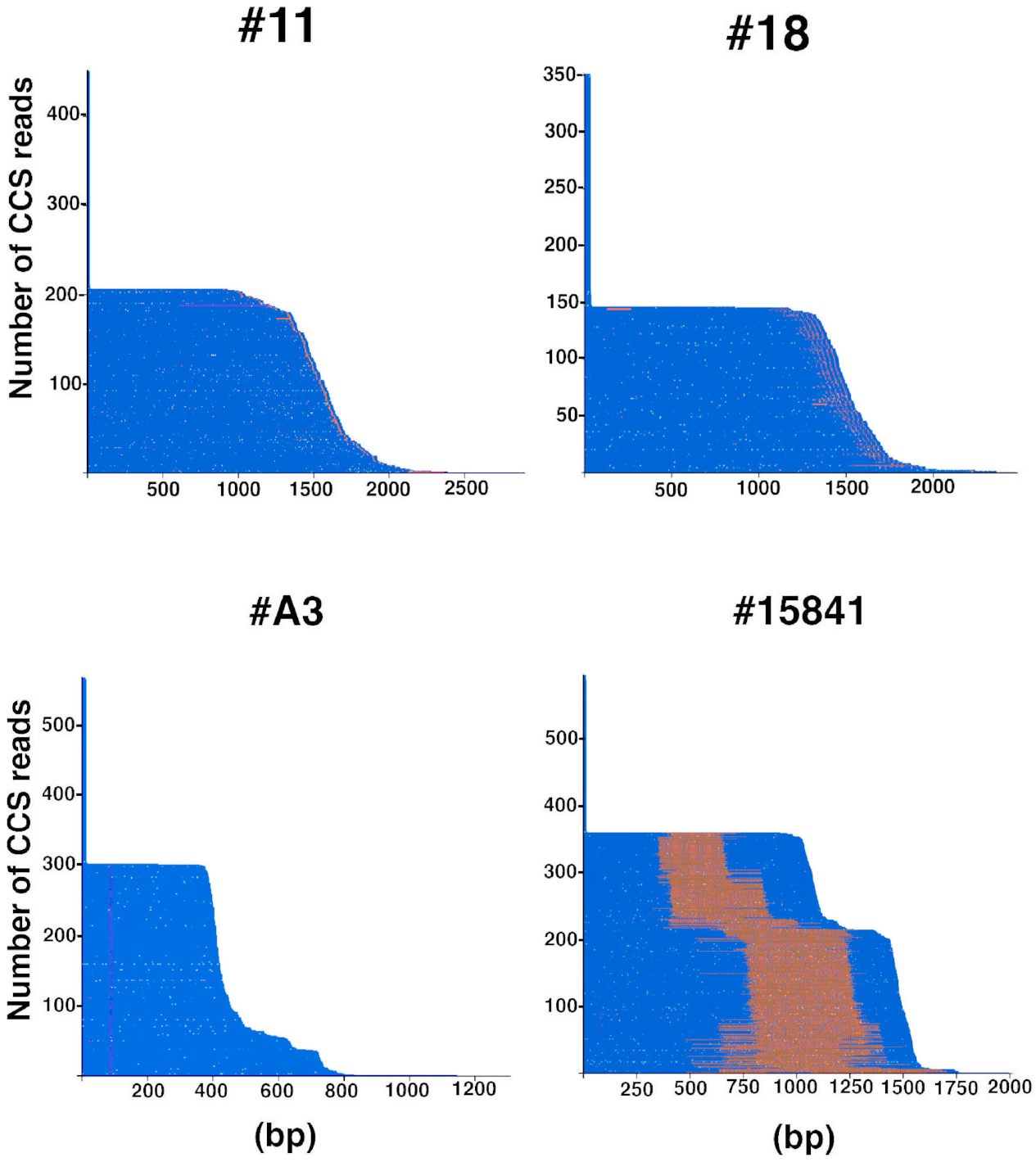
Waterfall plots outline the repeat structure of the normal and expanded alleles. Triplet repeat sequences are sorted from shortest to longest and each individual molecule is represented by a series of colored dots on a horizontal line. Each dot represents a single repeat unit. The CTG repeat is represented in blue whereas the CCG repeat is represented in orange. The highest peaks represent the normal allele. The x-axis shows the size of the repeat in base pair (bp).

### Identification of DM1 families carrying CCG-enriched expanded allele

DM1 patients in families 1 and 2 were identified at the neuro-myology department (Myology Institute Paris) as atypical DM1 patients in whom mild or no muscle symptoms were reported. In addition, no cardiac and severe respiratory abnormalities were observed (Figure 5 and Table 5). In particular, individual #14 is completely asymptomatic even after 12 years of follow up. Individual #14 has neither motor weakness nor myotonia, even on electromyography. Respiratory and cardiac investigations remain normal. These clinical aspects differ from what is usually observed in DM1 patients with similar age and CTG repeat size (ranging from 360 to 1000 by Southern blotting)^4^. To better characterize the *DMPK* mutation in DM1 members of these two families, targeted sequencing without amplification was performed in individuals #1, #14, #16 and #17 where conventional PCR failed. The amplification free DM1 locus enrichment is shown in table 6 where only 8 expanded allele HiFi reads were obtained in sample #1, making it difficult to estimate repeat size and somatic mosaicism in this patient (Table 6). The estimated size of the repeats in samples #14, #16, and #17 was 299, 224 and 360 CNG repeats, respectively, which is lower than what was estimated by Southern blot (360, 430 and 1000 CNG repeats, respectively) (Table 7). These results may be explained by an insufficient number of HiFi reads obtained for each sample, particularly for samples #16 and #17. Strikingly, large expanded alleles enriched in CCG triplet repeats were identified, which explains the difficulties in amplifying the CNG repeat expansions in these patients (Figure 5). Individual #14 mutated alleles (CTG)61-71(CTGCCG)115(CTG)1-5 are composed of more than 35% CCG repeat interruptions. The expanded allele structure observed in patient #14 differs from the expanded allele structure observed in her sister #1. The presence of more than 35% hexamer (CTGCCG) appears to stabilize the repeat in the blood of patient #14 (aged 35 years, Figure 6). In family 2, patients #16 and #17 carry an expanded allele with more than 85% CCG interruptions. The major allele structure observed in #16 is (CTG)7(CCG)(CTG)3(CCG)197(CTG)16 and differs from that observed in her daughter #17, where the major allele structure is (CTG)7(CCG) (CTG)3(CCG)330(CTG)19. No-Amp sequencing results showed a maternal CNG repeat expansion between individuals #16 and #17, suggesting that DM1 expanded allele with block CCG interruptions is unstable across generations in this family. Furthermore, somatic mosaicism was observed in these two patients with two different distributions of CTG repeat number (Figure 6). The low number of HiFi reads did not allow an accurate estimate of somatic mosaicism in these patients (Table 6). However, for the first time, a new variant at the DM1 locus composed of more than 85% CCG repeats was identified using the No-Amp targeted sequencing method.

**Table 5:** Clinical features in DM1 families 1 and 2.

|  | **Age sample**<br>**(Years)** | **Age at onset**<br>**(Years)** | **CTG repeat**<br>**number**<br>**(Southern Blot)** | **Weakness** | **Myotonia** | **Cardiac**<br>**defect** | **Respiratory**<br>**defect** |
| --- | --- | --- | --- | --- | --- | --- | --- |
| **#1** | 45 | 36 | N/A | Axial, proximal, distal and<br>facial | Yes (36y) | No | Vital capacity<br>91% and<br>Hypercapnia |
| **#14** | 35 | Asymptomatic | 360 | No | No | No | No |
| **#16** | 48 | 38 | 430 | Very mild facial | No | No | No |
| **#17** | 21 | 20 | 1000 | Distal weakness of the<br>hand<br>(Hirayama disease) | ~15y | No | No |

**Table 6:** HiFi reads in DM1 patients of families 1 and 2.

| **Individuals** | **#1** | **#14** | **#16** | **#17** |
| --- | --- | --- | --- | --- |
| **Hifi reads** | 926 | 652 | 510 | 1916 |
| **DM1 locus** | 158 | 192 | 127 | 471 |
| **Control loci** | 393 | 230 | 220 | 1275 |
| **Expanded allele** | 8 | 79 | 42 | 38 |
| **Non-Expanded allele** | 150 | 113 | 85 | 433 |
| **% reads on-target** | 59,61 | 64,88 | 64,12 | 91,13 |
| **% reads off-target** | 40,39 | 35,12 | 35,88 | 8,87 |

**Table 7:** CTG repeat size and interruptions in DM1 patients of families 1 and 2.

| **Individuals** | **#1** | **#14** | **#16** | **#17** |
| --- | --- | --- | --- | --- |
| **Median Expanded allele (CNG)** | N/A | 299 | 224 | 360 |
| **Interruptions** | 52 CCG | 115 CCG | 198 CCG | 331 CCG |
| **% CCG repeats** | N/A | 39% | 88% | 92% |
| **CNG repeat size (Diag)** | N/A | 360 | 430 | 1000 |

**Figure 5:**
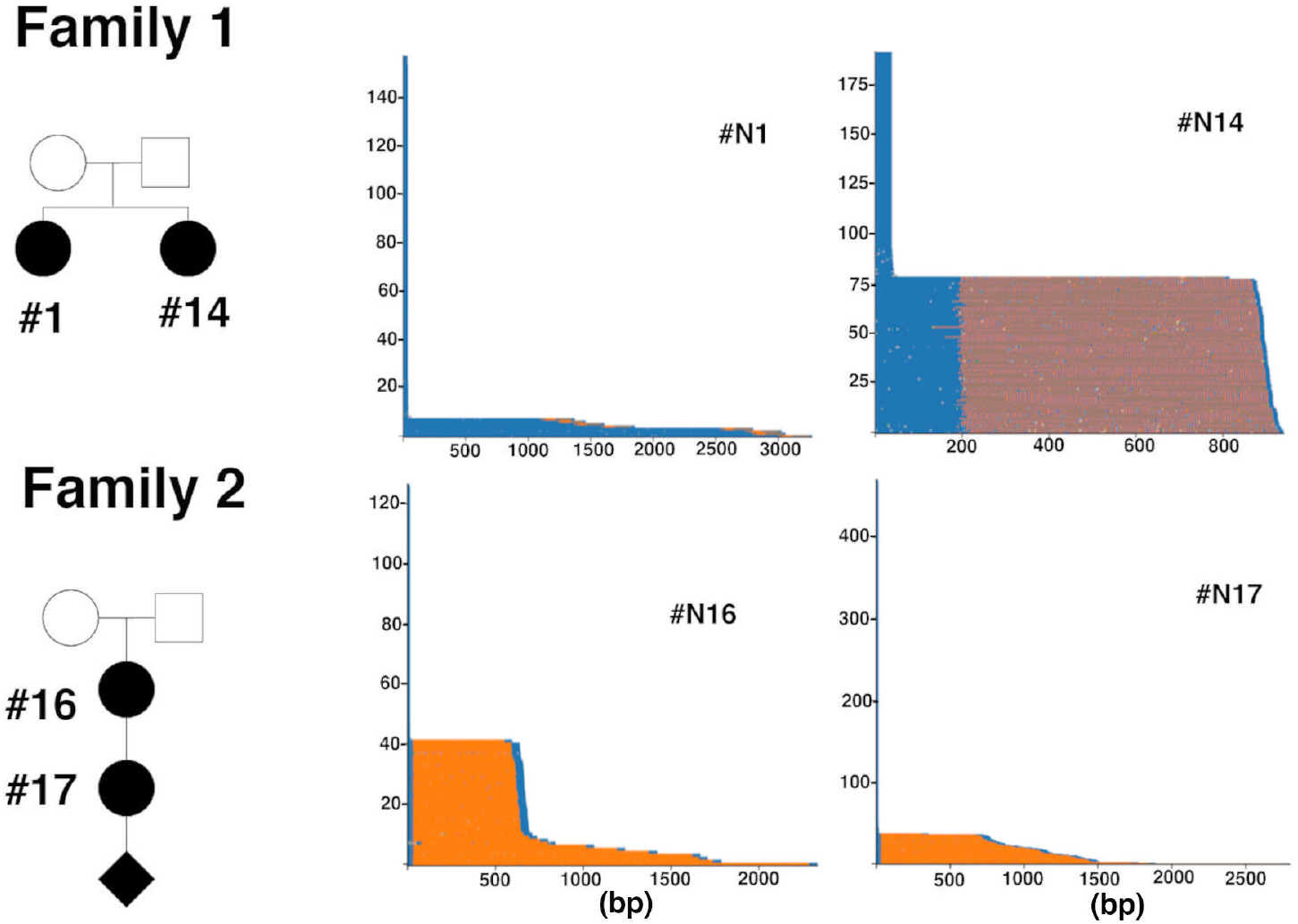
Waterfall plots outline the repeat structure of the normal and expanded alleles in DM1 families 1 and 2 (PCR failed). Triplet repeat sequences are sorted from shortest to longest and each individual molecule is represented by a series of colored dots on a horizontal line. Each dot represents a single repeat unit. The CTG repeat is represented in blue whereas the CCG repeat is represented in orange. The highest peaks represent the normal allele. The x-axis shows the size of the repeat in base pair (bp).

**Figure 6:**
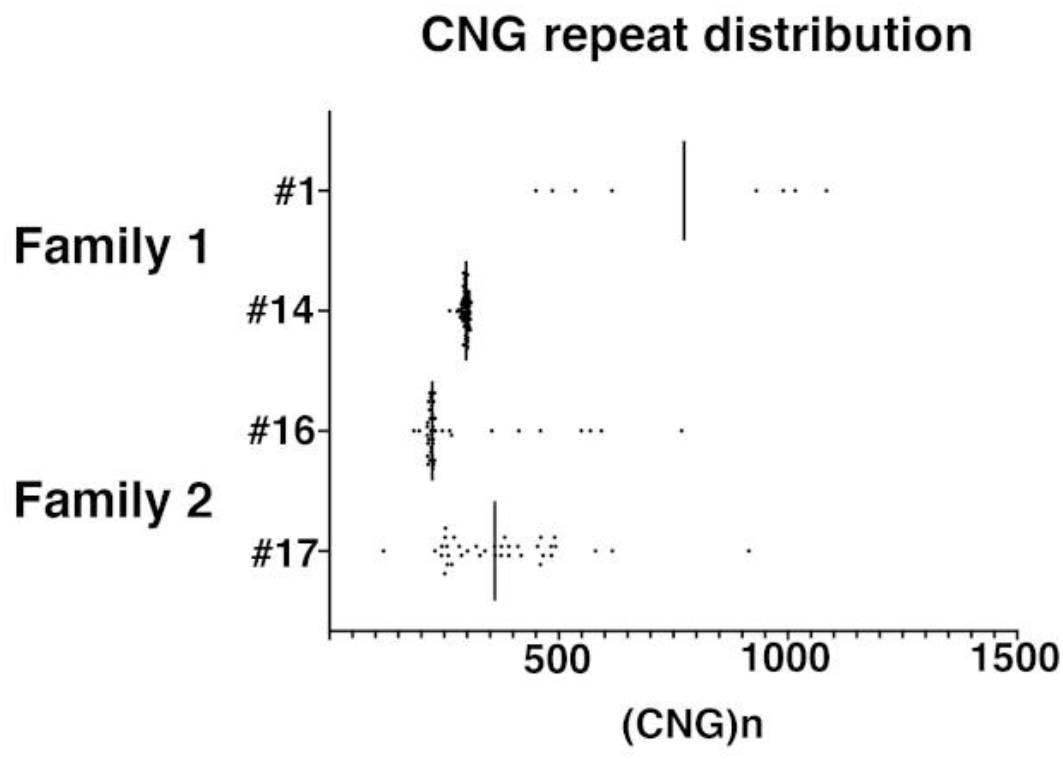
Distribution of the size of the repeat in families 1 and 2. The x-axis shows the CTG repeat number. Each dot represents the CTG repeat size of one HiFi read. The black line represents the median. 8-79 single molecule HiFi reads of expanded allele were analyzed in the No-Amp experiment.

## Discussion

In DM1, current routine molecular approaches do not provide simultaneous precise characterization of the size and the sequence of the DM1 mutation with precision as well as the dynamics of repeat instability in DM1 patients (Table1). These parameters are important DM1 disease modifiers^3^. In some cases, incomplete genetic analyses may lead to an inaccurate prognosis in DM1 patients. New genetic tests for DM1 are becoming essential to solidify the diagnosis, aid prognosis and improve care with shorter time frames. Today, the diagnostic delays remain too long for DM1 families who are often diagnosed at the birth of their child with a severe form of the pathology. Recently, we have shown that PacBio long-read sequencing from amplicons can sequence large CTG repeat expansions in DM1 patients and detect CTG repeat interruptions^27^. Nevertheless, this approach remains limited by PCR amplification. Here, we demonstrated the PacBio No-Amp targeted sequencing method is effective to sequence through previously inaccessible DM1 locus (large expanded alleles and GC-enriched interrupted alleles). In the DM1 population, more than 30% of patients carry large or interrupted expanded alleles^4,10^. This number is certainly underestimated by the fact that the size and sequence of the CTG repeat expansion is not systematically determined during diagnosis. Amplification-free targeted sequencing allowed highly accurate sequence determination of hundreds of expanded DNA molecules in several DM1 samples.

We have observed up to 50% DM1 on-target rate in HiFi sequencing reads when targeting only the DM1 locus with the Amplification-free targeted sequencing method. The on-target rate is affected by the quality of genomic DNA where high molecular weight DNAs provide the highest no-amp sequencing yield in this study (Table 3). Recently, it has also shown that high molecular weight DNA, yielding DNA ranging from 50 to 300+ kilobases in length, generates the highest quality long-read sequencing data of complex metagenomes^31^. The amplification-free targeted sequencing method is also sensitive to the origin of DNA samples. Variable results with DNA samples extracted from different tissues or cell types have been observed (data not shown). In order to continue improving the methodology in the future, additional studies to understand how different DNA sources and extraction methods are affecting this targeted sequencing application are needed.

During library preparation without amplification, a distribution close to 50%-50% of normal and mutated sequencing reads was observed in most sequencing data. However, an overrepresentation of the normal allele in patients with the largest expansion was observed. This phenomenon can be explained by lower HiFi read conversion rate because of the sequencing read length limitation and lower immobilization efficiency of DNA-polymerase complex on the Sequel sequencing chip. Long DNA molecules also have higher probability to carry damages susceptible to exonuclease degradation during sequencing library preparation. The combination of these factors could lead to non 50%-50% distribution of sequencing reads on normal and expanded alleles. Although including circular consensus (CCS) reads with predicted read quality below QV20 (accuracy <99%) in data analysis could recover additional reads with long repeat expansion, the option was not explored in this study to strengthen the identification of CNG interruptions. Improved long template immobilization efficiency and better sequencing read length on the newer SMRT sequencing platform could help overcome these limitations and provide more even coverages between normal and expanded alleles.

Removal of the PCR amplification step improves the accuracy of the measurement of the inherited CTG repeat expansion as well as the measurement of somatic mosaicism observed in blood or immortalized cells in most DM1 patients. Using No-Amplification targeted sequencing, we were also able to accurately detect interruptions and determine their number in blood and/or immortalized cells of all DM1 patients, regardless of the number of reads.

For the first time, DM1 patients with an expansion composed of more than 85% CCG associated with mild or no muscle symptoms and the absence of cardiac and severe respiratory abnormalities were identified by this method. This finding is consistent with recent studies showing that interrupted expanded alleles are associated with a decrease of the severity of DM1 symptoms^10,11,17,19–21^. It also raises question about systematic analysis of interruptions in DM1 patients. Altogether, our results confirm the need to revisit molecular diagnosis with No-Amp long-read sequencing to improve genotype-phenotype correlations and thus genetic counseling and prognosis in DM1.

Interestingly, heterogeneity in the sequence of the expansion, characterized by different expansion lengths and varying number of repeat interruptions, was detected in families 1 and 2. The heterogeneity in the number and type of interruptions observed in expanded interrupted alleles across generations (family 2) and in the sibling blood of family 1 suggests instability of the interruptions despite the fact that the same interruptions can stabilize the pure CTG repeat expansion^13–18,22^. The emergence of new interruptions in the expansion may result from multiple processes including DNA polymerase errors during replication, error-prone DNA repair or recombination. This method opens new avenues to understand the role of interruptions in DM1. The different mechanisms involved in the formation of CNG interruptions and instability of CTG repeats at the DM1 locus remain unknown and need to be identified in patients carrying the interrupted expanded allele using new models.

In conclusion, this method allows to simultaneously obtain high resolution information on the number of repeats, a complete and accurate sequence and a measure of somatic mosaicism even for long repeats in the same assay. This approach is highly relevant in DM1, which is one of the most complex trinucleotide disorders associated with the largest expansion and with high genetic and clinical variability. No-amplification targeted sequencing gives us the opportunity to better understand the dynamics of CTG repeat instability and genotype-phenotype association in DM1. This method will be extremely useful to validate innovative therapeutic strategies aiming at decreasing the CTG.CAG repeat length and thus stop the progression of the disease in DM1 patients.

## Acknowledgements

The authors would like to thank the DM1 patients and colleagues at the Myology Centre for Research, Curie Institute and PacBio for helpful discussions and comments. We specially thank Sonia Lameiras, Sylvain Baulande, Tina Alaeitabar, Lori Aro, Deborah Moine and David Stucki for their active involvement in this project. This work was supported by the Institut National de la Santé et de la Recherche Médicale, Sorbonne Université, Institut de Myologie (Accelerator project) and the 2019 Targeted Sequencing SMRT Grant.

## References

1. Khristich AN, Mirkin SM. On the wrong DNA track: Molecular mechanisms of repeat-mediated genome instability. Journal of Biological Chemistry, 2020, 295:4134–70

2. Harper PS. Myotonic dystrophy. London, W.B. Saunders, 2001

3. Tomé S, Gourdon G. DM1 Phenotype Variability and Triplet Repeat Instability: Challenges in the Development of New Therapies. IJMS, 2020, 21:457

4. De Antonio M, Dogan C, Hamroun D, Mati M, Zerrouki S, Eymard B, Katsahian S, Bassez G. Unravelling the myotonic dystrophy type 1 clinical spectrum: A systematic registry-based study with implications for disease classification. Revue Neurologique, 2016, 172:572–80

5. Genet AJH. Review Article: Anticipation in Myotonic Dystrophy: New Light on an Old Problem Peter S. Harper, Helen G. Harley, William Reardon, and Duncan J. Shaw, n.d.: 7

6. Kamsteeg E-J, Kress W, Catalli C, Hertz JM, Witsch-Baumgartner M, Buckley MF, van Engelen BGM, Schwartz M, Scheffer H. Best practice guidelines and recommendations on the molecular diagnosis of myotonic dystrophy types 1 and 2. Eur J Hum Genet, 2012, 20:1203–8

7. Savić Pavićević D, Miladinović J, Brkušanin M, Šviković S, Djurica S, Brajušković G, Romac S. Molecular Genetics and Genetic Testing in Myotonic Dystrophy Type 1. BioMed Research International, 2013, 2013:1–13

8. Morales F, Couto JM, Higham CF, Hogg G, Cuenca P, Braida C, Wilson RH, Adam B, del Valle G, Brian R, Sittenfeld M, Ashizawa T, Wilcox A, Wilcox DE, Monckton DG. Somatic instability of the expanded CTG triplet repeat in myotonic dystrophy type 1 is a heritable quantitative trait and modifier of disease severity. Human Molecular Genetics, 2012, 21:3558–67

9. Morales F, Vásquez M, Corrales E, Vindas-Smith R, Santamaría-Ulloa C, Zhang B, Sirito M, Estecio MR, Krahe R, Monckton DG. Longitudinal increases in somatic mosaicism of the expanded CTG repeat in myotonic dystrophy type 1 are associated with variation in age-at-onset. Human Molecular Genetics, 2020, 29:2496–507

10. Cumming SA, Jimenez-Moreno C, Okkersen K, Wenninger S, Daidj F, Hogarth F, Littleford R, Gorman G, Bassez G, Schoser B, Lochmüller H, van Engelen BGM, Monckton DG, on behalf of the OPTIMISTIC Consortium. Genetic determinants of disease severity in the myotonic dystrophy type 1 OPTIMISTIC cohort. Neurology, 2019, 93:e995–1009

11. Overend G, Légaré C, Mathieu J, Bouchard L, Gagnon C, Monckton DG. Allele length of the DMPK CTG repeat is a predictor of progressive myotonic dystrophy type 1 phenotypes. Human Molecular Genetics, 2019, 28:2245–54

12. Peric S, Pesovic J, Savic-Pavicevic D, Rakocevic Stojanovic V, Meola G. Molecular and Clinical Implications of Variant Repeats in Myotonic Dystrophy Type 1. IJMS, 2021, 23:354

13. Braida C, Stefanatos RKA, Adam B, Mahajan N, Smeets HJM, Niel F, Goizet C, Arveiler B, Koenig M, Lagier-Tourenne C, Mandel J-L, Faber CG, de Die-Smulders CEM, Spaans F, Monckton DG. Variant CCG and GGC repeats within the CTG expansion dramatically modify mutational dynamics and likely contribute toward unusual symptoms in some myotonic dystrophy type 1 patients. Human Molecular Genetics, 2010, 19:1399–412

14. Musova Z, Mazanec R, Krepelova A, Ehler E, Vales J, Jaklova R, Prochazka T, Koukal P, Marikova T, Kraus J, Havlovicova M, Sedlacek Z. Highly unstable sequence interruptions of the CTG repeat in the myotonic dystrophy gene. Am J Med Genet, 2009, 149A:1365–74

15. Botta A, Rossi G, Marcaurelio M, Fontana L, D’Apice MR, Brancati F, Massa R, G Monckton D, Sangiuolo F, Novelli G. Identification and characterization of 5’ CCG interruptions in complex DMPK expanded alleles. Eur J Hum Genet, 2017, 25:257–61

16. The Scottish Myotonic Dystrophy Consortium, Cumming SA, Hamilton MJ, Robb Y, Gregory H, McWilliam C, Cooper A, Adam B, McGhie J, Hamilton G, Herzyk P, Tschannen MR, Worthey E, Petty R, Ballantyne B, Warner J, Farrugia ME, Longman C, Monckton DG. De novo repeat interruptions are associated with reduced somatic instability and mild or absent clinical features in myotonic dystrophy type 1. Eur J Hum Genet, 2018, 26:1635–47

17. Santoro M, Masciullo M, Silvestri G, Novelli G, Botta A. Myotonic dystrophy type 1: role of CCG, CTC and CGG interruptions within DMPK alleles in the pathogenesis and molecular diagnosis: Variant interruptions in pathogenesis and molecular diagnosis of DM1. Clin Genet, 2017, 92:355–64

18. Tomé S, Dandelot E, Dogan C, Bertrand A, Geneviève D, Péréon Y, Simon M, Bonnefont J-P, Bassez G, Gourdon G. Unusual association of a unique CAG interruption in 5’ of DM1 CTG repeats with intergenerational contractions and low somatic mosaicism. Human Mutation, 2018, 39:970–82

19. Wenninger S, Cumming SA, Gutschmidt K, Okkersen K, Jimenez-Moreno AC, Daidj F, Lochmüller H, Hogarth F, Knoop H, Bassez G, Monckton DG, van Engelen Bgm, Schoser B. Associations Between Variant Repeat Interruptions and Clinical Outcomes in Myotonic Dystrophy Type 1. Neurol Genet, 2021, 7:e572

20. Miller JN, van der Plas E, Hamilton M, Koscik TR, Gutmann L, Cumming SA, Monckton DG, Nopoulos PC. Variant repeats within the DMPK CTG expansion protect function in myotonic dystrophy type 1. Neurol Genet, 2020, 6:e504

21. Ballester-Lopez A, Koehorst E, Almendrote M, Martínez-Piñeiro A, Lucente G, Linares-Pardo I, Núñez-Manchón J, Guanyabens N, Cano A, Lucia A, Overend G, Cumming SA, Monckton DG, Casadevall T, Isern I, Sánchez-Ojanguren J, Planas A, Rodríguez-Palmero A, Monlleó-Neila L, Pintos-Morell G, Ramos-Fransi A, Coll-Cantí J, Nogales-Gadea G. A DM1 family with interruptions associated with atypical symptoms and late onset but not with a milder phenotype. Human Mutation, 2020, 41:420–31

22. Pešović J, Perić S, Brkušanin M, Brajušković G, Rakočević-Stojanović V, Savić-Pavićević D. Repeat Interruptions Modify Age at Onset in Myotonic Dystrophy Type 1 by Stabilizing DMPK Expansions in Somatic Cells. Front Genet, 2018, 9:601

23. Morales F, Vásquez M, Santamaría C, Cuenca P, Corrales E, Monckton DG. A polymorphism in the MSH3 mismatch repair gene is associated with the levels of somatic instability of the expanded CTG repeat in the blood DNA of myotonic dystrophy type 1 patients. DNA Repair, 2016, 40:57–66

24. Flower M, Lomeikaite V, Ciosi M, Cumming S, Morales F, Lo K, Hensman Moss D, Jones L, Holmans P, Monckton DG, Tabrizi SJ, TRACK-HD Investigators, Kraus P, Hoffman R, Tobin A, Borowsky B, Keenan S, Whitlock KB, Queller S, Campbell C, Wang C, Langbehn D, Axelson E, Johnson H, Acharya T, Cash DM, Frost C, Jones R, Jurgens C, ‘t Hart EP, van der Grond J, Witjes-Ane M-NN, Roos RAC, Dumas EM, van den Bogaard SJA, et al. MSH3 modifies somatic instability and disease severity in Huntington’s and myotonic dystrophy type 1. Brain, 2019, 142:1876–86

25. Warner JP, Barron LH, Goudie D, Kelly K, Dow D, Fitzpatrick DR, Brock DJ. A general method for the detection of large CAG repeat expansions by fluorescent PCR. Journal of Medical Genetics, 1996, 33:1022–6

26. Monckton DG, Wong L-JC, Ashizawa T, Caskey CT. Somatic mosaicism, germline expansions, germline reversions and intergenerational reductions in myotonic dystrophy males: small pool PCR analyses. Human Molecular Genetics, 1995, 4:1–8

27. Mangin A, de Pontual L, Tsai Y-C, Monteil L, Nizon M, Boisseau P, Mercier S, Ziegle J, Harting J, Heiner C, Gourdon G, Tomé S. Robust Detection of Somatic Mosaicism and Repeat Interruptions by Long-Read Targeted Sequencing in Myotonic Dystrophy Type 1. IJMS, 2021, 22:2616

28. Robinson JT, Thorvaldsdóttir H, Winckler W, Guttman M, Lander ES, Getz G, Mesirov JP. Integrative genomics viewer. Nat Biotechnol, 2011, 29:24–6

29. Gomes-Pereira M, Bidichandani SI, Monckton DG. Analysis of Unstable Triplet Repeats Using Small-Pool Polymerase Chain Reaction. Trinucleotide Repeat Protocols, vol. 277, New Jersey, Humana Press, 2004, pp. 061–76

30. Tomé S, Nicole A, Gomes-Pereira M, Gourdon G. Non-Radioactive Detection of Trinucleotide Repeat Size Variability. PLoS Curr, 2014. https://doi.org/10.1371/currents.md.ad50113b899fa1352ce70c087eead706

31. Florian Trigodet, Lolans Karen, Fogarty Emily, Shaiber Alon, Morrison Hilary, Barreiro Luis, Jabri bana, Eren A Murat. High molecular weight DNA extraction strategies for long-read sequencing of complex metagenomes. Molecular Ecology Resources, n.d.

